# The Brightening Sky and the Bug’s Advance: Unraveling the Drivers of Oxycarenus lavaterae Range Expansion

**DOI:** 10.1101/2025.09.07.674691

**Authors:** Volodymyr Tytar

## Abstract

Analyzing the Lime Seed Bug’s (*Oxycarenus lavaterae*) European range expansion, optimized Maxent models and comprehensive occurrence data (2007-2025) reveal a swift northward and eastward spread, with a distinct “rapid expansion” phase starting in 2017. Key drivers include minimum and maximum temperatures, and importantly, downward shortwave radiation (DSR). Increased DSR, linked to “global brightening” from reduced air pollution since the 1990s, provides crucial thermal benefits. This enables the bug’s basking behavior to effectively elevate body temperatures, mitigating cold stress and enhancing overwintering survival in newly colonized northern regions. Focusing on Ukraine and Latvia, optimal habitat is predicted in Ukrainian regions like Transcarpathia, aligning with observed occurrences, while Latvia shows minimal invasion chances. This study rigorously integrates statistical modeling (including SHAP analysis) with biological insights, demonstrating how temperature extremes and DSR act as physiological “bottlenecks” for the species’ successful adaptation and expansion. The findings advance understanding of insect range dynamics under climate change and regional air quality improvements, providing critical insights for biodiversity conservation and targeted pest management. Furthermore, the presented methodologies facilitate citizen science efforts for ongoing ecological monitoring, empowering broader community participation in tracking environmental responses. Continued interdisciplinary research on these climatic and anthropogenic factors is vital for refining predictive models and informing adaptive management in a changing world.

## 1. Introduction

Climate change is undeniably reshaping the distribution of species across the globe. The Sixth Assessment Report of the Intergovernmental Panel on Climate Change (IPCC) highlights that average planetary temperatures have risen by nearly 1.2°C over the last century due to human-driven warming, with projections indicating continued increases (Lee et al., 2023). As temperature and precipitation patterns alter (Forster et al., 2024), many species are being compelled to shift their geographical ranges to find suitable environmental conditions (Bellard et al., 2011; Harvey et al., 2023; Zurell et al., 2024). This phenomenon is primarily driven by the need for species to track their fundamental climate niche – the specific set of primarily temperature and moisture conditions they are adapted to survive and reproduce within (Tingley et al., 2009). However, while range shifts often involve species tracking their pre-existing environmental niches, they can also in particular cases create opportunities for novel adaptations to arise, but whether these range shifts are a novel adaptation is a nuanced question.

In response to climate change, poleward and altitudinal migration is a commonly observed phenomenon among species. As temperature and moisture conditions shift, many species are expanding their ranges towards higher latitudes and elevations to track suitable environmental conditions. Latitudinal range shifts have especially been documented in the Northern Hemisphere, and chiefly between 30° and 60° latitudes (Lenoir et al., 2024). These shifts have profound implications for biodiversity conservation and ecosystem functioning, highlighting the urgent need to understand and mitigate the impacts of climate change on the natural world.

True bugs, belonging to the suborder Heteroptera, represent a small fraction (around 3%) of the non-native arthropods that have established themselves in Europe (Roques et al., 2010). Despite their relatively in the meantime low numbers, they pose a significant risk as potential pests, causing damage to crops and natural vegetation, or becoming a nuisance in indoor urban environments during the colder winter months.

In this paper we focus on one species, the Lime Seed Bug, *Oxycarenus lavaterae* Fabricius 1787 (Hemiptera: Lygaeidae) (Fig. 1), which already occurs in Europe with a low economic impact and has the potential to spread naturally, though passive dispersal by human-mediated translocations is assumed a probable cause of spread of this species, for instance in Bulgaria (Simov et al., 2012). Therefore, the status of this insect as a non-native species in Europe is not entirely straightforward. While its recent spread and establishment in many parts of the continent suggest an invasion, the species has historically been present in the warm Palaearctic region of the Mediterranean Basin (Péricart, 2001). Therefore, its expansion could also be seen as a range extension from its native southern European distribution rather than a strict introduction of a truly alien species as, for instance, the Colorado Potato Beetle (Weber, 2003). However, it is not known whether this is a consequence of global warming or an adaptation of the species to colder conditions than in its area of origin, or perhaps both.

**Fig. 1.**
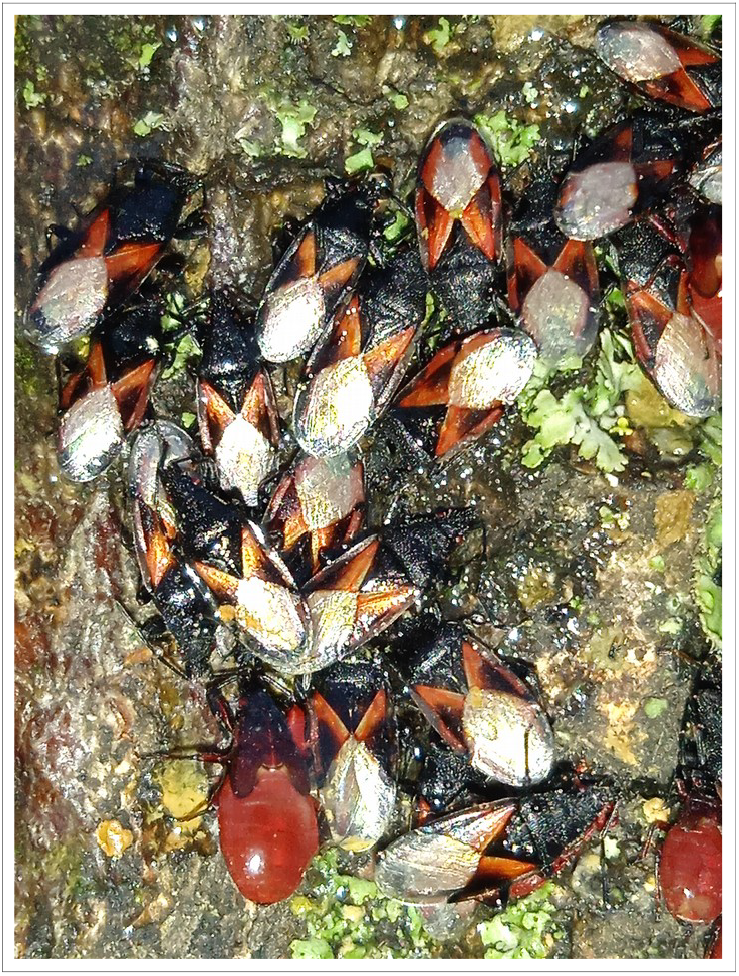
*Oxycarenus lavaterae*, 22.11.2024, Ukraine, Kyivska oblast, Bila Tserkva raion, 49.742344°N, 30.145343°E, by Serhij Oksenenko; UkrBIN

On the European continent, this species has been reported in the following years and countries: 1994 in Hungary; 1995 in Slovakia and Spain; 1996 in Serbia; 1998 in Bulgaria and Bosnia and Herzegovina; 1999 in France; 2001 in Austria; 2002 in Switzerland; 2003 in Finland; 2004 in Germany; Czech Republic and Croatia; 2007 in Netherlands and England; 2009 in Romania, Slovenia, and Greece; 2014 in Poland and Belgium; 2017 in Turkey; 2018 in Macedonia; 2019 in Luxemburg; 2020 in Russia (Bunescu et al., 2023; EPPO Global Database, 2010, https://gd.eppo.int/reporting/article-310). Since 2015 the species has been discovered in Ukraine (UkrBIN, https://ukrbin.com/index.php?id=347116%2F%29.&lang=1; single record, Korolevo in Transcarpathia). Numerous first records across Europe in recent decades and the clear pattern of northward and eastward movement strongly indicate that *O. lavaterae* is indeed a species undergoing a significant expansion of its home range within Europe (https://www.forestpests.eu/).

This study employs distribution and climate data for *O. lavaterae*, applying an optimized Maxent model to predict the species’ distribution (Phillips et al., 2006). We modelled the species’ niche models using standard procedures (Sillero, Barbosa, 2021) with the algorithm implemented in Maxent v3.4.4. In general, Maxent is a machine learning method using presence-only occurrence and background data, and compares the available environmental conditions in the study area (i.e., background) with the conditions used by the species, providing estimates of habitat suitability.

In parallel, modeling an advancing species like *O. lavaterae* in Maxent presents some unique challenges because the algorithm is primarily designed for modeling the current potential distribution of a species based on existing occurrence data and environmental variables. It doesn’t inherently model the process of invasion or its temporal dynamics. However, Maxent can be performed sequentially. The most important aspect is to have occurrence records of the expanding species collected at different time points as it spreads. Ideally, it would be to have data representing the initial occurrence points and subsequent spread over several time steps. Also relevant environmental data should ideally be available for the spatial extent of the considered study area and, if possible, for the different time periods. Analyzing the output of each Maxent model will give a snapshot of the species’ potential distribution at each time step. Unfortunately, in reality, occurrence data is often collected erratically, with varying effort and numerous gaps. The same largely applies to information gathered about the environment.

More often Maxent has been used to model the distribution and potential spread of invasive species, highlighting areas of expansion or contraction of suitable habitat. These studies use the body of occurrences collected to date in order to project current distributions into the future under different scenarios (in particular, climate change) (for example, Zhang et al., 2021; Mao et al., 2022; Liu et al., 2023; Adamu, Hussaini, 2024; Khwarahm, 2025).

In light of the aforementioned issues, taking *O. lavaterae* as the subject, we will be aiming to apply both a step-wise Maxent modeling approach using available time-series occurrence data to analyze the species’ spread over time, and a more common Maxent approach utilizing the entire occurrence dataset to project current and potential future distributions under different climatic scenarios. We are going to analyze the characteristics of its distribution patterns, predict its potential distributions, and evaluate the important environmental factors affecting its distribution. Based on the obtained findings, we aim to understand how relatively short- and long-term climate change, from the past to the future, affects the distribution patterns and ranges of the species, resulting in shifts in habitat distribution over time. Our intention is to make a special focus on Ukraine and Latvia as biogeographically contrasting areas, where Latvia is firmly within the northern, cooler, and more forested Boreal zone, while Ukraine acts as a transition zone with a wider range of climates and habitats, including the extensive steppes that are absent in Latvia (Roekaerts, 2002), not to mention Latvia’s greater distance from the likely source populations of the species under investigation.

## 2. Materials and Methods

### 2.1. Distributional Data

The geographic distribution data for *O. lavaterae* were compiled from two primary sources: (1) the Global Biodiversity Information Facility (GBIF.org, 2025), contributing a majority of the total records; and (2) the Ukrainian Biodiversity Information Network (https://ukrbin.com/). The initial dataset comprised 16,564 occurrence points, encompassing locations across Europe and North Africa.

To mitigate spatial sampling bias and potential geolocation errors we, after removing duplicate occurences, implemented a spatial thinning protocol performed in SAGA GIS using the ‘Points Thinning’ module (Conrad et al., 2015). As a rule of thumb, data points should be at least 2-3 cells apart in order to reduce autocorrelation (https://damariszurell.github.io/EEC-MGC/index.html). Following M. A. Nuñez and K. A. Medley (Nuñez, Medley, 2011), we measured the spatial autocorrelation of occurrences by calculating Moran’s I for multiple distance classes using the GeoDa software (Anselin, Koschinsky, 2022); values <0.3 were considered acceptable for building meaningful SDMs (Lichstein et al., 2002). Eventually, 3,134 occurence points were left for analysis. The longitude and latitude coordinates (WGS84 datum), and years of collection (ranging from 2007 up to 2024) of the sample were stored in an Excel database and converted into CSV format for the establishment of the Maxent model.

### 2.2. Environmental Variables and Processing

#### 2.2.1. Environmental Data Sources

SDMs are primarily climate-driven, meaning that the variables used to develop them typically portray climatic factors (Kriticos, 2012).

For the step-wise approach the TerraClimate global database, which has a spatial resolution of 2.5 arc-minutes or 5 km^2^ at the equator, serves as the source for the extraction of monthly climate and climatic water balance variables spanning from 1958 to the present (Abatzoglou et al., 2018). This dataset provides a detailed, long-term record of essential climate variables and derived water balance indices, making it valuable for ecological and hydrological studies. Available terraclimate variables can be downloaded using the ‘TerraclimateR’ package (https://julianselke.github.io/TerraclimateR/) and clipped to the region of interest (i.e., Europe and North Africa) by employing the ‘terra’ package (https://cran.r-project.org/web/packages/terra/index.html). At large, the potential geographical distribution of species can be inferred from maximum temperature, minimum temperature, relative humidity, rainfall, and other environmental factors (Wiens et al., 2009), therefore we have centered our attention on maximum and minimum temperatures, precipitation and downward shortwave radiation (DSR) at the surface, representing the total amount of solar radiation that reaches a horizontal unit area at the Earth’s surface. DSR is an important part of the Earth’s energy balance, driving Earth’s system’s energy, water, and carbon cycles (Liang et al., 2019). In our case, it is important to note that DSR has an indirect but significant influence on variables of several climate datasets (for instance, WorldClim, https://www.worldclim.org/) commonly used in correlative species-distribution modelling because it is a primary driver of both temperature and, to some extent, precipitation patterns. Using the PAST v.2.17c software package (Hammer et al., 2001), monthly variables were averaged to annual.

For building Maxent models of the current distribution and potential spread of the bug species we employed a 1 km global dataset of historical (1979–2013) and future (2020–2100) Köppen–Geiger climate classification and 12 bioclimatic variables (Table 1), resampled to a spatial resolution of 2.5 arc-minutes. These provide detailed descriptions of annual averages, seasonality, and stressful conditions of climates (Cui et al., 2021). The dataset referred to as KGClim is publicly available via http://glass.umd.edu/KGClim. The dataset presents six historical 30-year periods of the observational record and four future 30-year periods under four Representative Concentration Pathways (RCPs). For our purposes we downloaded data for the historical period of 1988-2017 and future (2030) under the RCP 4.5 scenario, often described as a low to moderate emissions scenario (Thomson et al., 2011) or semi-optimistic (Zeraatkar et al., 2025).

**Table 1.** List of bioclimatic variables derived from downscaled monthly climate data of the KGClim dataset.

| Bioclimatic variables & description |
| --- |
| BIO1 Annual mean temperature ( ° C) |
| BIO2 Temperature of the warmest month ( ° C) |
| BIO3 Temperature of the coldest month ( ° C) |
| BIO4 Annual precipitation (mm) |
| BIO5 Precipitation of the warmest half year (mm) |
| BIO6 Precipitation of the coldest half year (mm) |
| BIO7 Precipitation of the driest month (mm) |
| BIO8 Precipitation of the driest month in the warmest half year (mm) |
| BIO9 Precipitation of the driest month in the coldest half year (mm) |
| BIO10 Precipitation of the wettest month (mm) |
| BIO11 Precipitation of the wettest month in the warmest half year (mm) |
| BIO12 Precipitation of the wettest month in the coldest half year (mm) |

### 2.3. Modelling approach

MaxEnt was configured to conduct 10 cross-validate replicate runs with 10,000 random background points. Consequently, the average of the 10 predictions from all replicates was used for our analysis. In addition, 80% of occurrence data were selected for model training and 20% for model testing. To determine the optimal model complexity, we explored all combinations of the regularization parameter from 0.5 to 4 at intervals of 0.5 and potential combinations of four feature classes: linear, quadratic, product and hinge, using the ‘gridSearch’ function in the ‘SDMtune’ R package (Vignali et al., 2020) and assessing model selection by the area under the curve (AUC) of the Receiver Operating Characteristic (ROC) plots (Hanley, McNeil, 1982), based on both the training and validation datasets. Commonly used approaches recommend removing correlated predictor variables before modeling to avoid multicollinearity, which affects model projections (Zhao et al., 2022). There are several statistical packages offering functions that reduce collinearity in predictors, however in our work they were not employed because the benefits of using all available variables may outweigh the drawbacks of collinearity. Latest research indicates that modelling with correlated climate variables increases accuracy of predictions (Hanberry, 2023). Moreover, complex models such as Maxent take advantage of existing collinearity in finding the best set of parameters (De Marco, Nóbrega, 2018).

The model’s predictive accuracy was measured using the widely recognized AUC statistic. AUC scores range from 0 to 1, with values closer to 1 reflecting strong discriminatory power in distinguishing habitat suitability for the considered species (Wang, 2007). A score of 0.5 indicates random prediction, while values above this threshold demonstrate increasing reliability for differentiating between probable presence and absence zones (Phillips et al., 2017). Because AUC remains a controversial criterion (Lobo, 2008), for greater confidence we employed the continuous Boyce index, CBI (Boyce et al., 2002), one of the most reliable presence-only evaluation metrics. It is continuous and varies between −1 and +1. Positive values indicate a model that presents predictions that are consistent with the distribution of presences in the evaluation dataset, values close to zero mean that the model is not different from a random model (Hirzel et al., 2006). Estimates of the CBI were reached using an R code (https://github.com/jmrmcode/contboyceindex.git; author Juan M. Requena-Mullor) and classifying habitats as suitable or unsuitable for the survival of the target species based on a decision cut-off point equaling the Maximum training sensitivity plus specificity logistic threshold, as described by Liu et al. (2005).

### 2.4. Spatiotemporal issues

As climate change progresses, species are shifting their ranges to find suitable habitats. This often involves moving towards or away from the equator. In this respect, ‘Distance to Null Island’ (D2NI) can be a useful factor in analyzing species distribution patterns. Null Island is the point where the equator intersects the prime meridian (0° latitude, 0° longitude) in the Atlantic Ocean and D2NI is simply the distance from any point on Earth to this specific location calculated by using Pythagoras’ theorem (Şevgin, Öztürk, 2024). By monitoring changes in the D2NI of locations where a species is found, researchers can assess how climate change is affecting their shifting distribution and potentially predict future ranges. In order to track these shifts, the aforementioned Excel database composed of columns “longitude”, “latitude” and “year of collection” was supplemented with a column representing the D2NI.

As this dataset represents an 18 years long time series of D2NI values, subsequent analyses might benefit from smoothing techniques to reduce noise, performing trend analysis to identify long-term patterns, and searching for breakpoints that indicate abrupt changes and/or turning points (Hamilton, 2020). In this context corresponding modules of the PAST software have been used. We explored trends over time with the Mann-Kendall test and Sen’s slope, through the R package ‘trend’ (Pohlert, 2023). The Mann-Kendall test is a non-parametric test for monotonic trend detection in a time series. It analyses differences in signs between two consecutive dates: if a trend is present, the sign values will tend to increase or decrease, constantly. Sen’s slope is a non-parametric estimator of the magnitude of a linear trend in a time series. It provides a measure of the average rate of change in the data over time. For finding breakpoints the ‘BreakoutDetection’ R package was employed, implementing a technique for robustly, i.e., in the presence of anomalies, detecting single or multiple change points in univariate time series (https://github.com/twitter/BreakoutDetection).

### 2.5. Conditioning factors

1. The important environmental factors influencing the potentially suitable habitats of *O. lavaterae* were searched using the percentage and permutation contributions and Jackknife test results produced by the Maxent software. Here we prioritize permutation importance as far as it provides a more stable and interpretable measure of a variable’s influence on the model’s ability to accurately predict species distribution (Zeraatkar et al., 2025). To further explore the impact of these identified environmental factors, we employ a SHAP framework from XAI (i.e., eXplainable artificial intelligence) to rank and uncover the most influential drivers (Lundberg, Lee, 2017; Farooq et al., 2022). SHAP (SHapley Additive exPlanations) is a unified framework in explainable artificial intelligence used to interpret the output of any machine learning model by assigning each feature an importance value for a particular prediction. We post-processed the best model results with SHAP by comparing what a model predicts with and without the predictor for all possible combinations of predictors at every single observation. The predictors are then ranked according to their contribution for each observation and averaged across observations. Another useful item are dependence plots. In our case, the R package ‘shap-values’ (https://github.com/pablo14/; author Pablo Casas) in a modified version was used to perform the SHAP analysis. Also the package can produce SHAP dependence plots, a model agnostic visualization tool that helps understand the relationship between a variable and the model’s prediction (Niemann et al., 2020).

The application of SHAP for understanding the influence of environmental factors on species distribution has, until today, seen limited exploration, but is now being investigated more widely (for instance, Song, Estes, 2023).

## 3. Results & Discussion

### 3.1. Trends regarding D2NI

Descriptive statistics of D2NI such as mean, maximum and minimum, difference between them, were computed on an annual scale for the study period (Table 1).

The availability and quantity of observational species occurrence records have greatly increased due to technological advancements and the rise of online portals, such as the Global Biodiversity Information Facility (GBIF), however it is well-established that such records are biased in time and space (Petersen et al., 2021). Therefore, we tested the need for a smoothing technique to reveal underlying trends and patterns likely obscured by the noise and inconsistencies in the data collection process using the original and the 3-point moving average smoothed data, graphically shown in Fig. 2 and presented in Table 2. Even so, we understand that smoothing can result in the loss of some, maybe important, information (Berthouex, Brown, 2002).

**Table 1.** A statistical summary of distances between Null Island (0° latitude, 0° longitude) and occurrence locations of *Oxycarenus lavaterae* recorded throughout the years 2007-2024.

| year | maxD | minD | diffD | meanD | year | maxD | minD | diffD | meanD |
| --- | --- | --- | --- | --- | --- | --- | --- | --- | --- |
| 2007 | 48.83 | 37.59 | 11.24 | 42.59 | 2016 | 52.46 | 36.89 | 15.57 | 43.76 |
| 2008 | 52.26 | 37.30 | 14.96 | 40.94 | 2017 | 52.89 | 32.78 | 20.11 | 45.60 |
| 2009 | 51.55 | 36.73 | 14.83 | 42.63 | 2018 | 53.79 | 31.94 | 21.85 | 47.03 |
| 2010 | 48.64 | 33.02 | 15.63 | 40.04 | 2019 | 54.90 | 33.00 | 21.90 | 49.98 |
| 2011 | 47.28 | 32.02 | 15.27 | 39.78 | 2020 | 57.00 | 32.84 | 24.16 | 50.23 |
| 2012 | 47.29 | 33.61 | 13.67 | 40.61 | 2021 | 56.33 | 31.93 | 24.39 | 48.10 |
| 2013 | 51.29 | 33.74 | 17.54 | 41.26 | 2022 | 60.15 | 31.83 | 28.32 | 49.61 |
| 2014 | 51.29 | 33.74 | 17.54 | 41.26 | 2023 | 60.15 | 31.73 | 28.42 | 49.89 |
| 2015 | 50.13 | 32.00 | 18.13 | 41.17 | 2024 | 60.93 | 32.12 | 28.81 | 50.78 |
| abbreviations: maxD - maximum value of D2NI; minD - minimum value of D2NI; diffD - difference between maxD and minD; meanD - arithmetic mean of D2NI values |  |  |  |  |  |  |  |  |  |

**Table 2.**
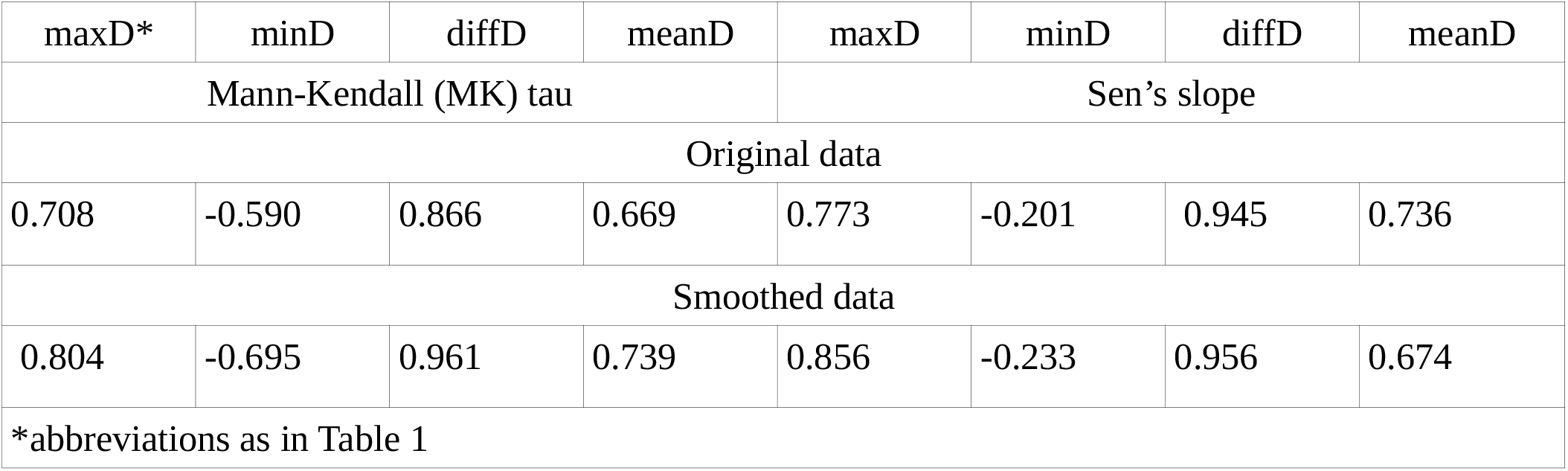
The Mann-Kendall (MK) and Sen’s slope estimation applied to detect trends in annual D2NI series (n=18) recorded during the study period; p<0.05.

| maxD* | minD | diffD | meanD | maxD | minD | diffD | meanD |
| --- | --- | --- | --- | --- | --- | --- | --- |
| Mann-Kendall (MK) tau |  |  |  | Sen's slope |  |  |  |
| Original data |  |  |  |  |  |  |  |
| 0.708 | -0.590 | 0.866 | 0.669 | 0.773 | -0.201 | 0.945 | 0.736 |
| Smoothed data |  |  |  |  |  |  |  |
| 0.804 | -0.695 | 0.961 | 0.739 | 0.856 | -0.233 | 0.956 | 0.674 |
| *abbreviations as in Table 1 |  |  |  |  |  |  |  |

**Fig. 2.**
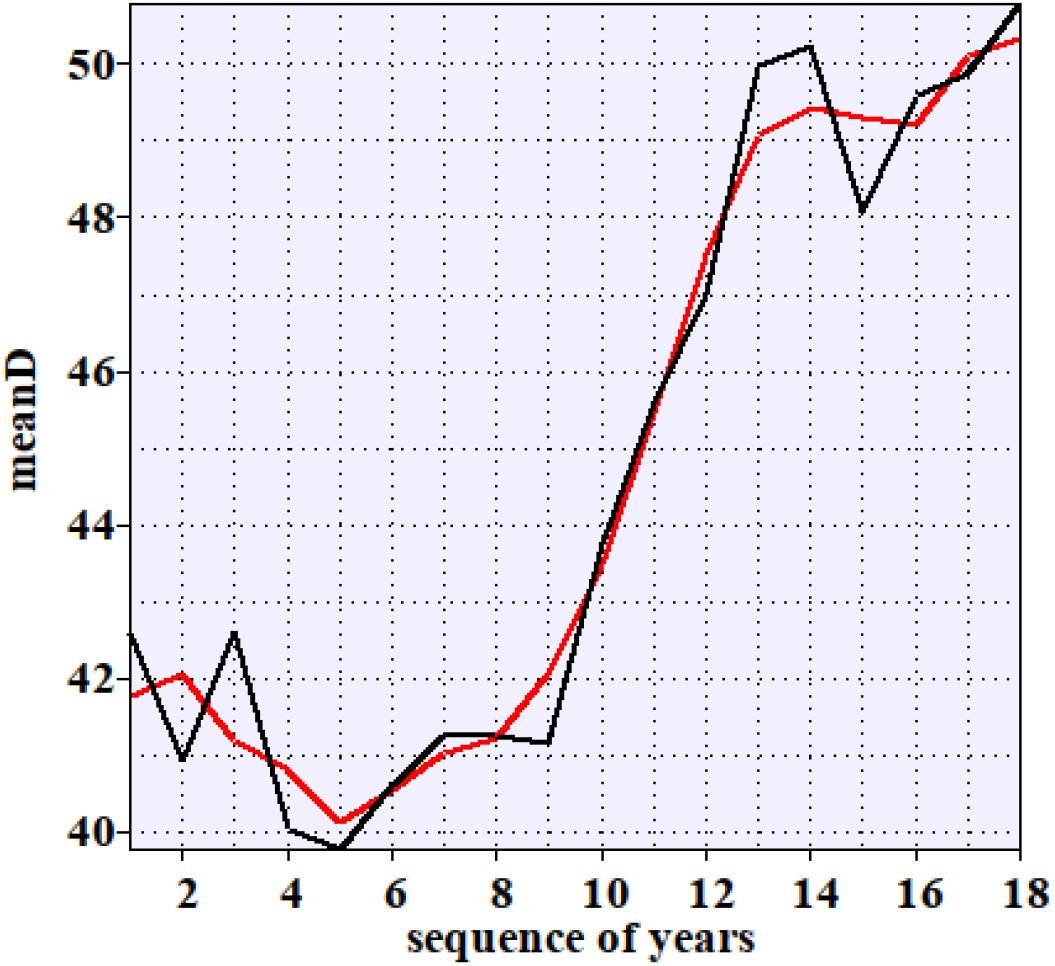
Annual trend for the mean D2NI over the 18 year time period: black line - original values of the arithmetic mean of D2NI values, red line - the 3-point moving average smoothed data; graph drawn using PAST ver. 2.17c

In general, despite possible variations in sampling locations and intensity each year, both original and smoothed data for *O. lavaterae* showed similar trends, as evidenced by the figures in Table 2, and we have chosen to primarily base our reasoning and conclusions on the original data. For example, figures representing temporal changes regarding the difference between the maximum and minimum values of D2NI in both instances indicate a very steep upward trend with a significant rate of change over time. Basically, this reflects the rapid expansion of the species in recent years and can serve as a quantification of this process. Instead of just saying the expansion of an invasive species is happening, we now have a specific number that describes its rate and strength (as indicated by Sen’s slope and tau).

Considering the overall consistency between the original and smoothed data in reflecting the general trends of *O. lavaterae* regarding changes over time, we can look into the specific behaviors of other aspects of the D2NI trend issues.

Firstly, examination of the temporal evolution of the D2NI’s minimum values shows a pronounced negative downward trend characterized by a significant decreasing rate of change over the 18 year time period, whereas maximum values show the opposite. This most likely suggests a dual expansion of the species, both towards the equator and northwards; however, the northward shift has occurred at a considerably greater rate, nearly four times the absolute value of the rate of the southward expansion. Secondly, considering mean values, the overall trend exhibits a strong positive increase of D2NI values over the study period with a noticeably high rate of change.

While the average D2NI values show a strong upward trend over the entire study, this increase doesn’t appear consistent, as evidenced by the graph in Fig. 2. The initial period seems to lack significant change, followed by a much sharper rise later on. This non-uniform pattern suggests the potential presence of a breakpoint in the trend, warranting further investigation to identify when this shift occurred. Using the method = ‘amoc’ (At Most One Change) in the breakout() function from the BreakoutDetection library, we revealed a breakpoint at the 10-year mark of the observation period (with a p value <0.05, meaning a real change in the mean of our time series and not just random variation).

To analyze the trends before and after the identified breakpoint, smoothing the data might be necessary due to a seemingly insufficient number of data points to achieve statistical significance in the trend analysis for these shorter periods. Indeed, a trend analysis of the smoothed D2NI values within the time frame of 2007-2016 found no changes in the data over this specific period (p values substantially exceeding 0.05, suggesting no significant trend), whereas for the period of 2017 to 2024, we obtain a statistically sound (p < 0.05) output value of 0.929 (tau), which indicates a strong, monotonic increase in annual average D2NI values over this time period occurring at a noticeable rate of 0.542, as indicated by Sen’s slope. Speaking figuratively, the spread of *O. lavaterae* can be broadly divided into an earlier period of “staticity” up to 2016 and a subsequent phase of “rapid expansion” starting in 2017, however a closer look at the data presented in Fig. 1 suggests a deceleration in this later expansion phase. This visual indication of a slowdown aligns with the behavior often described by the Michaelis-Menten model, a nonlinear fitting option available in software like PAST, which is commonly used to model processes exhibiting an initial rapid increase followed by a plateau as limiting factors come into play. Indeed, the Michaelis-Menten model statistically confirms this observed deceleration and performs better, according to the Akaike information criterion, than other proposed in the PAST software models.

### 3.2. Model development and evaluation

As mentioned in the ‘Introduction’, in the first place of our analysis of the species’ spread over time we apply a Maxent modeling approach using available time-series occurrence points and relevant for the spatial extent of the study area environmental data encompassing the period from 2007 to 2024. Taking into account that this time frame is split, as we see from above, into two phases both the occurrences and the corresponding environmental variables have been accordingly assigned to two separate sets of data. Further we model the potential distribution of the species in two steps: before its pronounced home range expansion and after, giving a snapshot of the species’ potential distribution at each time step.

A total of 314 spatially filtered occurrence points of *O. lavaterae* from across its home range were used to model the potential distribution of the species that would accord to the time period between 2007 and 2016. As spelled out by the ‘SDMtune’ algorithm, optimal model complexity was reached by employing a regularization parameter of 0.5 and a combination of three feature classes: linear, quadratic, and hinge. This optimized model exhibited high AUC and CBI scores, 0.852±0.039 SD and 0.848±0.021 SD, respectively, making it sufficiently well-suited for projecting the habitat distribution of the considered species at this precise time interval.

Another set of 2,820 processed occurrence points of *O. lavaterae* was used to model the potential distribution of the species that would relate to the time period between 2017 and 2024. This set too represents the entire bug’s home range. Consistent with the previous findings using the ‘SDMtune’ algorithm, optimal model complexity was achieved by repeatedly applying a regularization parameter of 0.5 and the same combination of feature classes. AUC and CBI scores indicate a satisfactory result: 0.788±0.011 SD and 0.865±0.020 SD, respectively, with the CBI score being particularly noteworthy. Looking at these home rage models and clipping them to boundaries of Latvia and Ukraine allows using geostatistics available in SAGA GIS to assess how suitable has the environment been for the considered bug species at different time intervals. Corresponding habitat suitability scores based on the modeling exercises are presented in Table 3.

**Table 3.** Habitat suitability scores based on *Oxycarenus lavaterae* distribution models for different time periods.

| Time period | Habitat suitability scores |  |  |  |  |  |  |  |  |
| --- | --- | --- | --- | --- | --- | --- | --- | --- | --- |
|  | Europe & N. Africa |  |  | Latvia |  |  | Ukraine |  |  |
|  | Min.* | Max. | Mean | Min. | Max. | Mean | Min. | Max. | Mean |
| Models using predictors from the TerraClimate global database |  |  |  |  |  |  |  |  |  |
| 2007-2016 | 0.000 | 0.972 | 0.206 | 0.000 | 0.006 | 0.001 | 0.000 | 0.476 | 0.071 |
| 2017-2024 | 0.000 | 0.717 | 0.292 | 0.017 | 0.081 | 0.052 | 0.014 | 0.503 | 0.182 |
| Models using predictors from the KGCLim dataset |  |  |  |  |  |  |  |  |  |
| current | 0.000 | 0.924 | 0.135 | 0.013 | 0.224 | 0.064 | 0.000 | 0.371 | 0.090 |
| 2030 | 0.000 | 0.848 | 0.136 | 0.011 | 0.158 | 0.055 | 0.095 | 0.431 | 0.095 |

\*Min. - minimum, Max. - maximum, Mean - arithmetic mean of habitat suitability scores

Consistently across all considered instances, an upward trend in mean habitat suitability is apparent and proved to be statistically significant, as evidenced by the p-value <0.05, demonstrating a robust pattern. The very low scores for Latvia relating to both the period before 2017 and after are suggesting that conditions have not yet been established here that would favour the species’ presence. Indeed, at the moment there are no records of *O. lavaterae* from Latvia. The same largely applies to Ukraine for the time period of 2007-2016 during which a single finding of the bug species was made in 2015 in the Transcarpathian region close to the border with Hungary, where it is known since 1994. However, a real outburst of registered occurrences of the species came off in the period following 2017, with the number reaching 90. Undoubtedly, this increase may also be at least partially due to enhanced inventory efforts, but which mostly remain undocumented. Interestingly, the maximum habitat suitability score during the expansion period (0.717) is lower compared to the “static” period (0.972), implying that the species, particularly within the present-day northern expanses of its home range, is perhaps in some final stages of establishment, as suggested by the Michaelis-Menten model described above. It might not have thoroughly explored or adapted to the full range of suitable habitats available in the new region yet as it has in southern Spain, Near East and north Africa, areas regarded “native” (Péricart, 2001). Over time, the predicted suitability might increase as the species expands its realized niche and continues, though most likely at a slower pace, its northwards shift. The research team of O. Nedvěd et al. (2023), highlighting the critical role of increasing temperatures, posits that the species’ expansion is expected to continue if the mild winters that have prevailed in central Europe in recent years were to occur in more northern countries.

Having processed the home range models and clipped them to the boundaries of Latvia and Ukraine, the following step is to visualize the habitat suitability for the considered bug species, focusing on the example of Ukraine, and identify the regions within Ukraine that are predicted to be most susceptible to initial invasion. Next we overlay the species occurrence points within Ukraine onto a thresholded suitability map. In the first place we considered to choose the minimum training presence threshold (MTP), an approach preferred in modeling invasive species as the least stringent (Ouko et al., 2020; Baici, Bowman, 2023). However, given that the MTP threshold yielded overly inclusive predictions encompassing the entire country, we opted instead for the one percentile training presence logistic threshold to define suitable areas, where predicted areas with values below 0.12 were deemed unsuitable for *O. lavaterae*. The subsequent outcomes are displayed in Fig. 3.

**Fig. 3.**
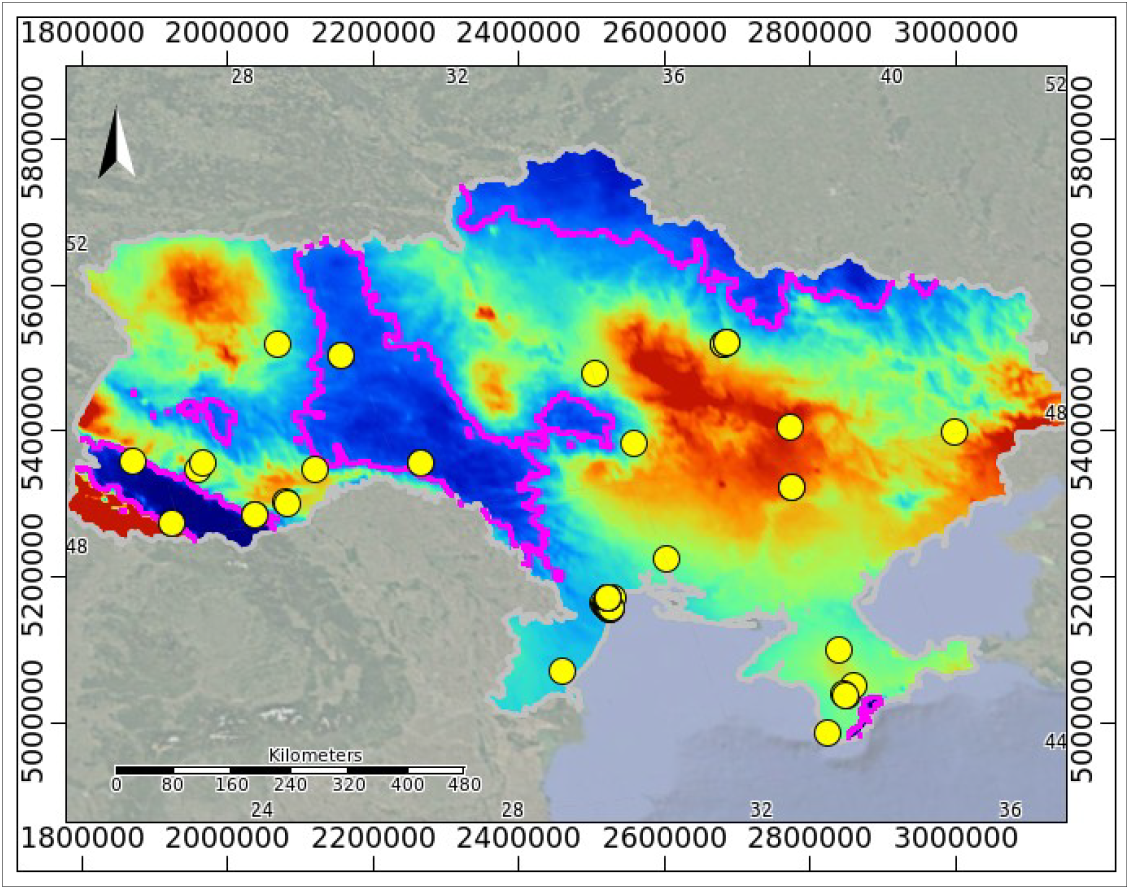
Habitat suitability (HS) map for *Oxycarenus lavaterae* in Ukraine based on occurrences for the time period of 2017-2024 and the TerraClimate dataset; colours show potential HS ranging from high (red) to low (navy blue); the fuchsia-coloured contour line represents the one percentile training presence logistic threshold; yellow circles symbolize point occurrences. Coordinate reference system: Lambert azimuthal equal-area projection.

A Chi-square test of homogeneity (or independence) was performed to compare the distribution of actual species occurrences with that of randomly generated presences. The highly significant Chi-square statistic (χ2=47.9, p<0.05) indicates that actual species occurrences are not randomly distributed. Instead, the species exhibits a strong preference or aggregation pattern that is highly unlikely to occur by chance.

The analysis above suggests that in terms of climate, best conditions in Ukraine for the invader are predicted to occur in the extreme south-west of the country within the administrative region of Transcarpathia (or Zakarpats’ka Oblast’), which borders Hungary, Slovakia and Romania from where first records of *O. lavaterae* were made between 1995 and 2009. Other areas include the Precarpathian region (primarily L’vivs’ka Oblast’), Chernivets’ka Oblast’, regions in the north-west (Volyns’ka and Rivnens’ka oblasts), oblasts located roughly in the center of the country (Dnipropetrovs’ka, Kirovohrads’ka, Poltavs’ka), and south-east (Zaporizs’ka, Donets’ka, Luhans’ka). Kyiv, the capital city, is too in this list, presumptively due to the phenomenon known as the urban heat island effect (Kokosha, 2024).

The following step in model development consisted in utilizing the entire occurrence dataset to project current and potential future distributions under different climatic scenarios by employing the KGClim dataset. In general, this is a widely adopted and common approach in ecological niche modeling involving an array of environmental predictors. We modeled situations for the historic period of 1984-2013 and the near-future period, including scenarios for 2020-2049 (2030s). Optimal model complexity was achieved by applying invariably a regularization parameter of 1.0 and a combination of four feature classes: linear, quadratic, product and hinge. The models showed reasonably high AUC and CBI scores across the analyzed periods: historic (1984-2013) with 0.817±0.006 SD and 0.866±0.021 SD, and 2030s with 0.817±0.010 SD and 0.855±0.022 SD. Corresponding habitat suitability scores based on the modeling exercises involving the whole home range and separately Latvia and Ukraine are presented in Table 3.

The Chi-square test performed in this case, which compared the distribution of actual species occurrences to that of randomly generated presences, yielded a lower though yet significant statistic (χ2=4.8, p<0.05). This result indicates that actual species occurrences are not randomly distributed; however, the observed preference or aggregation pattern is presumably weaker than that suggested by the TerraClimate model.

While no trends in mean habitat suitability were apparent in the considered cases (p>0.05), Latvia was an exception. However, even there, mean figures remained low (and are predicted to even diminish), suggesting a minimal chance for *O. lavaterae* to become widespread, whereas in Ukraine the invasion chances are above 50% higher. Due to the large similarity between current and future (2030) habitat suitability maps for Ukraine, only the current situation is presented (Fig. 4).

**Fig. 4.**
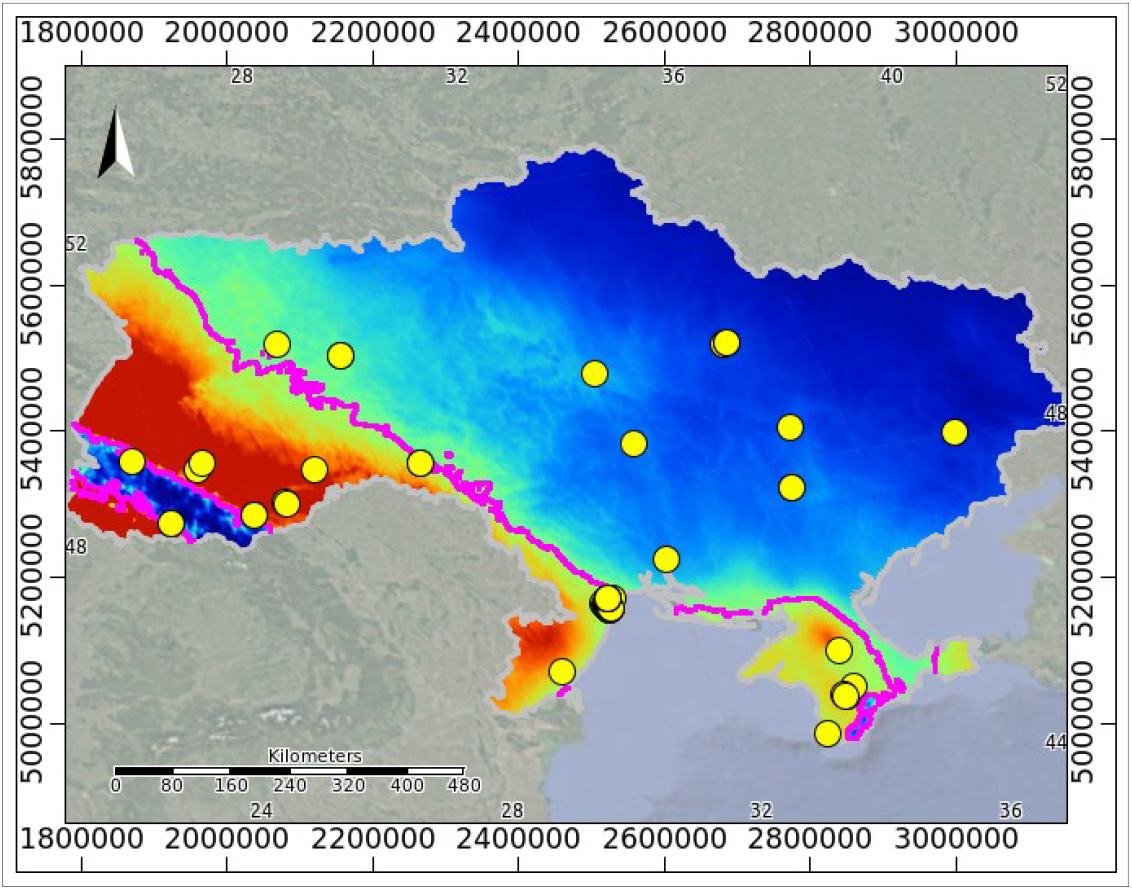
Current habitat suitability (HS) map for *Oxycarenus lavaterae* in Ukraine based on occurrences for the time period of up to 2025 and the KGClim dataset; colours show potential HS ranging from high (red) to low (navy blue); the fuchsia-coloured contour line represents the one percentile training presence logistic threshold; yellow circles symbolize point occurrences. Coordinate reference system: Lambert azimuthal equal-area projection.

Our analysis suggests that climatically, optimal conditions for the invader in Ukraine under the considered model are expected to occur in low-ground areas of Transcarpathia, primarily within L’vivs’ka, Ivano-Frankivs’ka and Chernivets’ka oblasts in the west of the country. In the south suitable areas are predicted to occur in Odesa Oblast’ and Crimea.

Given the detailed insights from the Maxent models regarding the Lime Seed Bug’s expansion and projected optimal climatic conditions across Ukraine, a weighted (by the corresponding AUC) average consensus model can now be developed to synthesize these findings into a more robust and comprehensive predictive map (Fig. 5). As far as earlier applied thresholds yielded overly inclusive predictions, we opted for the ten percentile training presence logistic threshold to define suitable areas, where predicted areas with values below 0.04 were considered unsuitable for *O. lavaterae*.

**Fig. 5.**
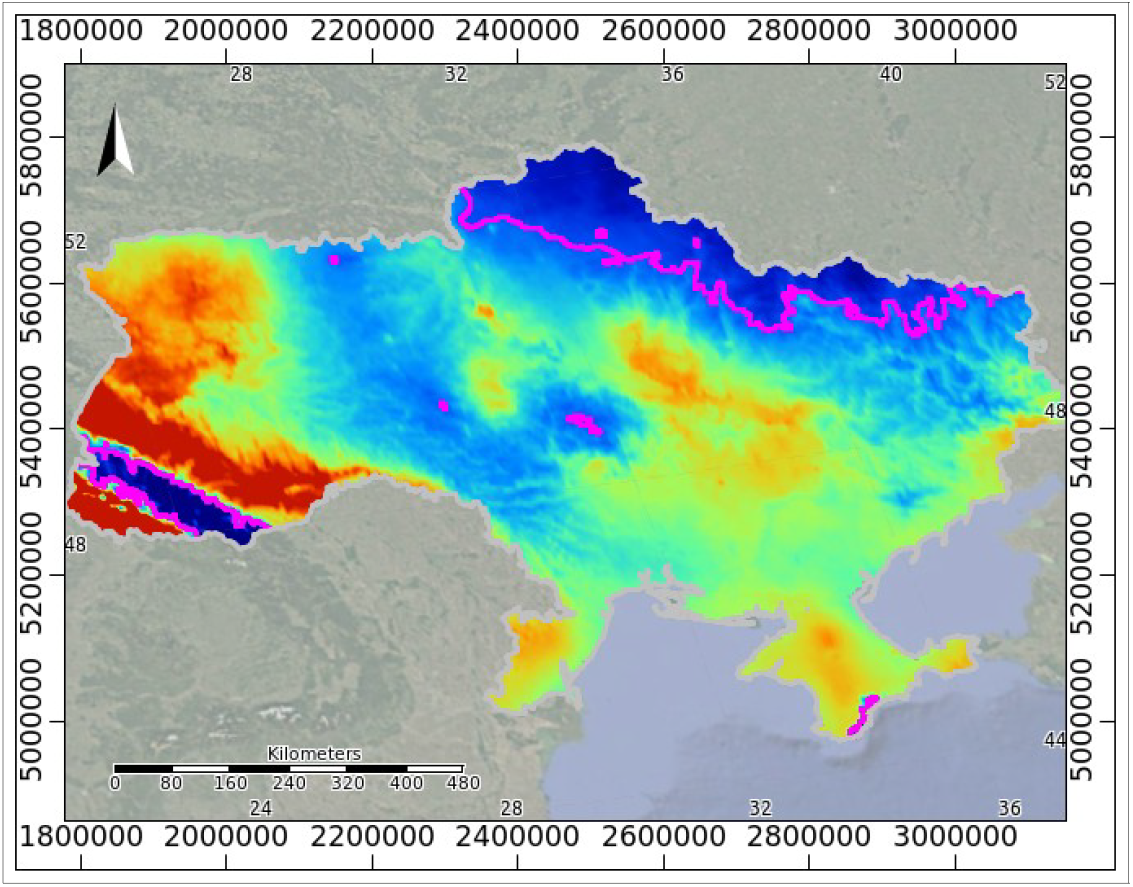
Current consensus habitat suitability (HS) map for *Oxycarenus lavaterae* in Ukraine; colours show potential HS ranging from high (red) to low (navy blue); the fuchsia-coloured contour line represents the ten percentile training presence logistic threshold. Coordinate reference system: Lambert azimuthal equal-area projection.

The weighted average consensus model allows for the ranking of Ukrainian regions (oblasts) by their average habitat suitability for *O. lavaterae*. Regions exhibiting above-average suitability, listed in descending order, include: L’vivs’ka, Transcarpathia, Chernivets’ka, Volyns’ka, Ternopil’s’ka, Ivano-Frankivs’ka, Crimea, Rivnens’ka, Dnipropetrovs’ka, Khersons’ka, Zaporizs’ka, and Odes’ka. As anticipated, the most favorable areas for the establishment of *O. lavaterae* are predominantly located in western Ukraine. Conversely, the northeastern regions appear least suitable. While the highlands of the Carpathians and Crimea are excluded from favorable habitats, it’s notable that the majority of the country possesses the potential to support the bug’s presence.

### 3.3. Importance of environmental factors

#### 3.3.1. TerraClimate models

The ecological presence of *O. lavaterae* within its Afro-European home range is shaped by a combination of environmental variables. By analyzing the permutation importance of individual variables used in the modeling process, we found that the environmental factors derived from the TerraClimate database contributing the most to the model showing the potential distribution of the species according to the time period between 2007 and 2016 were overwhelmingly represented by temperature- and energy-related features. Namely, minimum (46.4%) and maximum (25.6%) temperatures, and DSR at the surface (20.9%) were significant factors; annual precipitation made only a minor contribution (7.1%). The jackknife test of variable importance too clearly stressed the importance of minimum temperatures.

Following the species’ rapid expansion, temperature- and energy-related features remained highly significant, collectively contributing 86.3% to permutation importance. This period, however, saw a reordering of their individual influence. Specifically, maximum temperatures and DSR at the surface became more prominent, both almost equally increasing their contributions to 38.1% and 37%, respectively. Conversely, the importance of minimum temperatures was reduced to 13.8%, and annual precipitation made just a slightly larger contribution at 11.2%. Correspondingly, the jackknife test indicated that the maximum temperature factor appears to have the most useful information by itself, whereas DSR appears to have the most information that isn’t present in the other variables.

To further explore the impact of these environmental factors, we employed a SHAP framework (see section 2.5). For the time period 2007-2016 the minimum temperature factor upheld its significance, even increasing it up to 56.0%. Analysis of the corresponding SHAP dependence plot reveals that the impact of minimum temperatures on the model’s predictions follows an inverted U-shape curve. Habitat suitability shows a climbing trend as minimum temperatures increase, reaching a peak between the mean annual figures of approximately 12 to 13 degrees Celsius. Beyond this span, SHAP values quickly drop to zero and turn into negative figures, indicating a steep decline in habitat suitability as minimum temperatures become excessively high. This illustrates how both extremely low and extremely high minimum temperatures are detrimental, highlighting an optimal range for this environmental factor. In theory, a positive effect (i.e., SHAP values >0) on the model’s predictions occurs between 10.6 and 17.5 degrees Celsius. While the model’s predictions show a positive linear correlation with minimum temperatures (Pearson’s coefficient = 0.68, p <0.05) – suggesting generally better habitat suitability at locations with higher minimum temperatures – the SHAP analysis reveals a more nuanced, non-linear relationship: both extremely low and extremely high minimum temperatures are detrimental to habitat suitability, thus effectively captures the complexities of the environmental drivers beyond simple linear associations. This intricate relationship with temperature is directly tied to insect physiology as far as lower temperatures decrease metabolic activity, lead to reduced movement, feeding, and other physiological processes (Cossins, Bowler, 1987; Irlich et al., 2009; Lalouette et al., 2012). In this respect winter conditions, particularly the coldest months of December, January, and February, are decisive for mean annual minimum temperatures in Europe (Copernicus Global Climate Highlights Report, 2024). Conversely, higher temperatures increase metabolic rate and activity up to a certain point (Kingsolver et al., 2015). However, exceeding the optimal range, as seen in the SHAP analysis, leads to the worsening of habitat suitability. In that context there is a certain similarity with the Linden bug, *Pyrrhocoris apterus*, where winter warming was found to have a strong negative effect on overwinter survival (Rozsypal, 2024). Therefore, both excessively low temperatures, which slow down vital processes, and excessively high temperatures, which are likely to cause physiological damage, contribute to the observed decline in habitat suitability outside the optimal range.

For an exploration of the role of environmental factors in tailoring the model for the 2017-2024 period we repeatedly employed the SHAP framework. Contrary to the permutation importance pointed out in the Maxent model, SHAP showed the dominating significance of DSR (45.2%), with the maximum temperature factor (24.8%) ranking second. This insight from SHAP likely provides a more refined picture of individual factor contributions. In light of this, DSR requires our special attention. Similar to the relationship observed with ‘maximum temperature’, the SHAP dependence plot for DSR reveals an inverted U-shaped relationship with the model’s predictions, reaching a peak mean annual value of around 125 W/m^2^, which falls within a moderate range of 114-160 W/m^2^, typical for mid-latitudes of Europe (e.g., Germany, France, Poland, Ukraine). This emphasized role of DSR in shaping habitat suitability is particularly critical when considering the rapid northward spread of the lime seed bug across Europe. Originating from the south, this invasive insect has shown a remarkable expansion into mid-latitudinal regions, and recently even further north, a phenomenon that could be triggered by increasing DSR. While rising winter temperatures are a well-documented factor facilitating the overwintering survival of many thermophilic insects (Bale, Hayward, 2010; Robinet, Roques, 2010; Chen et al., 2011; Halsch et al., 2021), the availability of sufficient solar radiation during colder months perhaps plays an even more vital role. *O. lavaterae* adults are known to form large, dense aggregations on tree trunks, especially on the sun-exposed south or southwest sides of lime trees, as a key overwintering strategy (Bunescu et al., 2023; Ilea et al., 2023; Domagała, 2024). In these exposed microhabitats, direct DSR provides crucial thermal benefits, allowing the bugs to bask and elevate their body temperatures above ambient air temperatures, thereby mitigating the physiological stress of cold and reducing winter mortality.

This understanding sets the stage for two key observations. First, regression analysis reveals that in regions like Germany, heavily invaded by *O. lavaterae*, DSR has increased at a faster rate than maximum temperatures (p < 0.05). For this purpose we employed a linear regression model with an interaction term using the lm() function in the R statistical environment (R Core Team, 2020). The model allowed us to assess both the individual effects of year and data type, as well as their interaction, to determine if the rate of change over time differed significantly between DSR and maximum temperature. Second, this rise in DSR, roughly coinciding with the northward expansion of the lime seed bug, is likely driven by a decrease in air pollution leading to clearer skies. This is a very important and complex topic, often referred to as “global dimming” or “brightening” (Wild, 2010; 2012). From about the 1950s to the 1980s, many industrialized regions experienced a phenomenon known as “global dimming,” where the amount of DSR reaching the Earth’s surface significantly decreased. This was largely attributed to a rapid increase in anthropogenic aerosol emissions (especially sulfates) during that period. But since the 1990s, many regions (particularly in Europe and North America) have experienced a reversal, known as “global brightening.” This is primarily due to successful air pollution control policies that have significantly reduced the emission of reflective aerosols. As these aerosols are removed from the atmosphere, more DSR is able to reach the Earth’s surface, directly impacting microclimates (Yuan et al., 2021) and expectedly affecting insects that rely on solar basking for thermoregulation (like *O. lavaterae*). Notably, our breakpoint analysis also aligns with this observation. The increasing trends in DSR in newly colonized by the bug regions could therefore be a significant, yet often underestimated, driver of this species’ successful northward range expansion and overwintering capacity, enabling it to survive cold winters where it was previously restricted by freezing temperatures and limited solar energy availability. Its developed basking behavior now effectively taps into this crucial solar energy source, overcoming a key previous limitation. Interestingly, a comparable pattern is evident in wild populations of the linden bug, *Pyrrhocoris apterus*, however for this particular species winter basking behavior has been found to be rare (Rozsypal, 2024).

#### 3.3.1. KGClim models

Using permutation importance, our analysis of environmental factors for the years 1988-2017 from the KGClim database showed that temperature of the coldest month (27.4%) and annual precipitation (18.1%) were the most influential in predicting the species’ distribution. Together, these two factors explained almost half of the model’s variation. The jackknife test stressed exclusively the importance of temperature-related factors: the annual mean temperature and temperature of the warmest month. The significant contribution of the coldest month’s temperature in this case mirrors and underscores the importance of the minimum temperature factor disclosed in the earlier Terraclimate model analysis. Indeed, even though the species is capable of tolerating cold snaps reaching -10°C, prolonged exposure to winter temperatures below -15°C presents a severe threat, potentially wiping out as much as 99% of the hibernating population (Kalushkov et al., 2007; Nedvěd et al., 2014).

For our exploration of how the KGClim environmental factors shaped the model for the current period, we consistently used the SHAP framework focusing on the top five most important. SHAP revealed that temperatures of the warmest and coldest month were clearly the dominant factors, accounting each for 33.8 and 31.5% of the average absolute impact on the model’s predictions.

The SHAP dependence plot shows that the impact of temperatures of the warmest month on the model’s predictions forms a seen before inverted U-shaped curve, suggesting that there’s a more or less ideal range of warmest month temperatures for the habitat suitability, with values too low or too high being less favorable. Specifically, habitat suitability tends to increase as these temperatures rise, peaking roughly between 17 and 21 degrees Celsius. For temperatures of the coldest month, the SHAP dependence plot reveals a varying impact as temperatures increase. The graph sharply rises from -3.7 degrees Celsius, flattening out around 0 degrees. However, the impact then shows a steady increase again from 7.5 degrees, continuing up to 15.6 degrees. Interestingly, the cease of coordinated movement in *O. lavaterae* occurs at around -3.2 to -3.4°C (i.e., close to our -3.7°C), indicating the critical thermal minima (CT_min_) for the species (Käfer et al., 2020). We believe this finding effectively bridges statistical modeling (SHAP) with biological reality.

As for the future, permutation importance once more indicated the significant role of coldest month temperatures (26.2%) and of the precipitation of the wettest month in the warmest half-year (25.2%), together explaining more than half of the model’s variation. Differently, however, temperature-related factors – the annual mean temperature and the temperature of the coldest month – were exclusively emphasized by the jackknife test. Revisiting the SHAP analysis, the results highlighted the temperatures of the coldest and warmest months as the dominant drivers, contributing 33.9% and 33.2% respectively (together, over two-thirds) to the average absolute impact on the model’s predictions. Notably, this mirrors the dominant factors observed for the current period, suggesting a fundamental and robust influence of these climatic variables. The fact that both the permutation importance and SHAP analysis consistently highlight these temperature-related factors implies that the underlying relationships within our models are strong and stable across different scenarios or methods of assessment and they are likely identifying the most impactful levers in the system. Even as average temperatures rise, an organism’s survival or success might still be primarily determined by the absolute coldest temperature it experiences, or the highest temperature it can tolerate. These extremes act as physiological “bottlenecks” or “tipping points”, and their relative importance for a species’ fate will probably remain in place even as the overall climate shifts (Field et al., 2012).

## 4. Conclusions

A comprehensive analysis of the Lime Seed Bug, *O. lavaterae*, reveals its significant range expansion across Europe, with an examination of both past observations and future projections. Optimized Maxent models, utilizing time-series occurrence data from 2007 to 2024 and various environmental factors, clarify the drivers behind this spread. A “rapid expansion” phase, notably commencing in 2017, marks a key period in the species’ distribution dynamics.

*O. lavaterae* demonstrates a dual expansion pattern, moving towards both the equator and northwards, but the northward shift has occurred at a considerably faster rate. This expansion is closely tied to critical environmental factors, with minimum temperatures, maximum temperatures, and downward shortwave radiation (DSR) at the surface exerting overwhelming influence on habitat suitability. DSR, a key driver, appears to be a significant, yet often underestimated, factor facilitating the successful northward range expansion and enhanced overwintering capacity of the bug. This is partly due to the species’ developed basking behavior, allowing it to leverage increased DSR – likely a consequence of reduced air pollution and “global brightening” since the 1990s – for thermoregulation during colder months. This effectively overcomes a crucial prior limitation of freezing temperatures and limited solar energy availability.

Statistical modeling, including SHAP framework analysis, integrates with biological realities, demonstrating how temperature extremes and DSR function as critical physiological “bottlenecks” for *O. lavaterae*.

A weighted average consensus model effectively ranks Ukrainian oblasts by their average habitat suitability for O. lavaterae. Regions with above-average suitability are primarily concentrated in the west, while northeastern regions and the highlands of the Carpathians and Crimea are less suitable, however the model indicates that most of Ukraine has the potential to support the Lime Seed Bug’s presence. In contrast, Maxent models for Latvia show minimal invasion chances for the species.

Overall, this comprehensive analysis not only advances understanding of insect range dynamics in a changing climate but also provides critical insights for proactive biodiversity conservation and targeted pest management strategies amidst ongoing environmental shifts and varying regional impacts of climate change and air quality improvements. Furthermore, the methodologies and findings presented here are capable of facilitating citizen science efforts for ongoing monitoring, empowering broader participation in tracking species’ responses to environmental change. Continued exploration of the nuanced interplay of these climatic and anthropogenic factors will be vital to refine predictive models and inform adaptive management approaches.

